# Role of sleep quality in the acceleration of biological aging and its potential for preventive interaction on air pollution insults: findings from the UK Biobank cohort

**DOI:** 10.1101/2021.08.27.457922

**Authors:** Xu Gao, Ninghao Huang, Tao Huang

## Abstract

**Background:** Sleep has been associated with aging and relevant health outcomes, but their causal relationship remains inconclusive.

**Methods:** In this study, we investigated the associations of sleep behaviors with biological ages (BAs) among 363,886 middle and elderly-aged adults from UK Biobank. Sleep index (0 [worst]-6 [best]) of each participant was retrieved from six sleep behaviors: snoring, chronotype, daytime sleepiness, sleep duration, insomnia, and difficulties in getting up. Two BAs, the KDM-biological age and PhenoAge, were estimated by corresponding algorithms based on clinical traits, and their discrepancies with chronological age were defined as the age accelerations (AAs).

**Results:** We first observed negative associations between the sleep index and the two AAs, and demonstrated that the change of AAs could be the consequence of sleep quality using Mendelian randomization with genetic risk scores of sleep index and BAs. Particularly, one unit increase in sleep index was associated with 0.105- and 0.125-year decreases in KDM-biological age acceleration and PhenoAge acceleration, respectively. Furthermore, we observed significant independent and joint effects of sleep and air pollution (i.e. PM_2.5_ and NO_2_), another key driver of aging, on BAs. Sleep quality also showed modifying effect on the associations of elevated PM_2.5_ and NO_2_ levels with accelerated aging. For instance, an interquartile range increase in PM_2.5_ level was associated with 0.011-, 0.047-, and 0.078-year increase in PhenoAge acceleration among people with high (5-6), medium (3-4), and low (0-2) sleep index, respectively.

**Conclusions:** Our findings elucidate that better sleep quality could lessen accelerated biological aging resulting from exogenous exposures including air pollution.

**Funding:** Peking University Start-up Grant (BMU2021YJ044)

## Introduction

Aging is a gradual and progressive deterioration in biological system integrity, which is thought to arise from the accumulation changes at the cellular level [1]. It is accompanied by changes in sleep quality, quantity, and architecture, especially in elderly adults [2]. Nevertheless, the mutual causal association between sleep and accelerated aging is still in debate. Along with many physiological alterations in normal aging, sleep behaviors change with aging independent of many factors including medical comorbidity and medications [3]. Some researchers thus suggested that sleep disorders are the consequences of aging-related changes in neuroendocrine functions [3]. Meanwhile, sleep is also considered as a restorative process that not only allows for energy renewal but also for cellular restoration [2]. Hence, the other hypothesis is that the declined sleep quality may lead to accelerated aging by influencing the compensatory/resiliency systems of human body by triggering DNA damage and chronic inflammation [2]. The lack of an established accurate measurement of aging, however, hinders the scientists from elucidating the directionality and causal relationship between aging and sleep.

Aging is a sum of changes that occur at hierarchically organized levels in the human body [1], which makes it hard to be reflected by single age-related biomarkers employed in previous relevant studies. All individuals age chronologically at the same rate, but there is marked variation in their biological ages as we observe in real life that the people with the same chronological age may not share the same aging-related symptoms [1]. Some could experience age-related decline faster than others. Moreover, as biological aging is a complex biological process in multiple organ systems, a single aging-related biomarker, such as telomere length or oxidative stress biomarkers, may not be able to completely depict the whole landscape of the aging process of individuals due to the heterogeneity of cells [4]. Therefore, the identification of “biological age (BA)” has been proposed and been explored in the last 10 years. To date, several forms of BAs which could be estimated based on the functions of cardiovascular, metabolic, renal, immune, and pulmonary systems (e.g. KDM-biological age [5, 6] and PhenoAge [7]), or based on aging-related DNA methylation profiles (known as “epigenetic clock” [8]), have been developed. Their discrepancies with chronological age, i.e. the age accelerations (AAs), have been highly associated with aging-related health outcomes and mortality [8]. However, previous studies on the associations of sleep with BAs or aging related symptoms usually employed only one or two sleep behaviors or were conducted in a specific population with limited participants [9-13]. Therefore, there is a dearth of study exploring the associations between sleep and aging in larger populations with causal inference approaches to uncover their causal nature explicitly.

Additionally, air pollution, especially the fine particulate matter [PM <2.5 μm (PM_2.5_)], is a critical environmental exposure that could advance aging [14] and affect sleep quality [15]. Previous studies have linked aberrant accelerated epigenetic clocks with the increased exposure to various air pollutants in different populations [14]. Plenty of evidence has also documented increased risks of sleep disorders [15] and sleep-related neurological impairments, e.g. dementia and cognitive decline [16, 17], associated with elevated air pollution levels. Nevertheless, since the sleep-aging relationship is still dim, no studies yet evaluated the associations of air pollution, sleep, and aging simultaneously, which is paramount for developing interventions to mitigate the adverse effects of air pollution on aging or sleep quality.

Therefore, we examined the causal associations of sleep (as reflected by six sleep behaviors) with two BAs based on the measures of systemic aging in multiple organs among 0.36 million participants of UK Biobank, a national-wide population-based cohort study in the UK. We subsequently explored the associations of five major air pollutants (PM_2.5,_ PM with an aerodynamic diameter between 2.5 and 10 µm [PM_coarse_], PM with an aerodynamic diameter of less than 10 µm [PM_10_], nitrogen dioxide [NO_2_], and nitrogen oxides [NOx]) with BAs and sleep, and explored whether BAs could mediate or modify the associations of air pollution with sleep, or vice versa.

## Methods

### Study design and population

Study design and methods of UK Biobank have been reported in detail previously [18]. In brief, UK Biobank is a large-scale prospective study with 502,536 participants aged 37–73 years recruited in 2006–2010 with multiple follow-ups. At the initial visit, participants provided information on sleep and other health-related aspects through touch-screen questionnaires and physical measurements. Blood samples were collected for genotyping and biochemistry tests. UK Biobank research has approval from the North West Multicenter Research Ethical Committee. All participants provided written informed consent for the study. In this analysis, we included 363,886 participants with available data of sleep behaviors, measures of biological traits for BA construction, and air pollution. This report followed the Strengthening the Reporting of Observational Studies in Epidemiology (STROBE) reporting guideline.

### Assessment of sleep behaviors

We used six self-reported sleep behaviors in this study: snoring, chronotype, daytime sleepiness, sleep duration, insomnia, and difficulties in getting up in the morning. (1) Information on snoring was collected by asking ‘Does your partner or a close relative or friend complain about your snoring?’ with responses of (a) yes or (b) no; (2) Chronotype was assessed using the following question, ‘Do you consider yourself to be (a) definitely a “morning” person, (b) more a “morning” than “evening” person, (c) more an “evening” than “morning” person, or (d) definitely an “evening” person?’. (3) Daytime sleepiness was retrieved from the question ‘How likely are you to doze off or fall asleep during the daytime when you don’t mean to? (e.g. when working, reading, or driving)’ with responses of (a) never/rarely, (b) sometimes, (c) often, or (d) all of the time. (4) Sleep duration was retrieved from the reported hours of sleep by asking ‘About how many hours sleep do you get in every 24 h? (include naps)’. (5) Insomnia symptoms were obtained by asking ‘Do you have trouble falling asleep at night or do you wake up in the middle of the night?’ with responses of (a) never/rarely, (b) sometimes, or (c) usually. (6) Difficulty levels of getting up in the morning were assessed by asking ‘On an average day, how easy do you find getting up in the morning?’ with responses of (a) not at all easy, (b) not very easy, (c) fairly easy, or (d) very easy. According to the six sleep behaviors, we generated a sleep index. The low-risk categories of each component were no self-reported snoring, early chronotype (‘morning’ or ‘morning than evening’), no frequent daytime sleepiness (‘never/rarely’ or ‘sometimes’), normal sleep duration (7–8 h per day), reported never or rarely having insomnia symptoms, and getting up easy in morning (‘fairly easy’ or ‘very easy’). For each sleep behavior, the participant received a score of 1 if he or she was classified as the low-risk group or 0 if otherwise as the high-risk group. All component scores were summed to obtain a continuous sleep index ranging from 0 (worst) to 6 (best), with a higher index indicating a general better sleep quality. We further defined a sleep index category as high (5-6), medium (3-4), and low (0-2) based on the continuous sleep index.

### Assessment of biological ages

We computed the BAs derived from a total of 12 blood chemistry traits, systolic blood pressure, and lung function data (Table 1) with two commonly accepted algorithms, the Klemera-Doubal method (i.e. KDM) and the PhenoAge method. Both algorithms were initially trained in data from the National Health and Nutrition Examination Survey (NHANES) following the method originally described by Klemera et al. [5, 6] and Levine et al. [7] with two sets of nine clinical traits (Table 1). The selected traits employed in each algorithm and corresponding R code can be found in the R package ‘BioAge’ at: https://github.com/dayoonkwon/BioAge [19]. Missing values of each trait consisted of <10% of all traits and therefore were imputed by the median value of the corresponding trait. The residual differences between the estimated BAs and chronological age were considered as AAs. The residuals were calculated by a linear regression procedure in which one of the BAs was the outcome and chronological age was the independent variable.

### Exposure assessments

As previously described [20], the annual average concentrations of PM_2.5,_ PM_coarse_, PM_10_, NO_2_, and NOx were calculated centrally by the UK Biobank using a Land Use Regression model developed by the ESCAPE project. Land Use Regression models calculate the spatial variation of annual average air pollutant concentration at participants’ home addresses given at the baseline visit, using the predictor variables obtained from the Geographic Information System such as traffic, land use, and topography. The exposure data of PM_2.5_, PM_coarse_, and NOx were collected in 2010, while annual concentrations of NO_2_ and PM_10_ were available for several years (2005, 2006, 2007, and 2010 for NO_2_ and 2007 and 2010 for PM_10_). Averaged values of NO_2_ and PM_10_ were included in the analysis.

### Measurements of covariates

We included age, sex, body mass index (BMI), race, smoking status, healthy alcohol intake status, healthy physical activity status, years of education (<10 years), and prevalent hypertension and diabetes as potential confounders. Height and weight were measured by trained nurses during the baseline assessment center visit, and BMI was calculated by dividing weight in kilograms by the square of height in meters. Healthy alcohol intake status was defined as: male: <28g/day; female: <14g/day. Healthy physical activity status was defined as: ≥150 min/week moderate or ≥75 min/week vigorous or 150 min/week mixed (moderate + vigorous) activity. The Metabolic Equivalent Task (MET) minutes based on items from the short International Physical Activity Questionnaire (IPAQ) was adopted to assess physical activity. The history of hypertension and diabetes was based on self-reported information and medical records.

### Construction of genetic risk scores of sleep index and biological ages

Detailed information about genotyping, imputation, and quality control in the UK Biobank study has been described previously [18]. We created two genetic risk scores (GRSs) for sleep index and BAs/AAs for the MR analyses, respectively. A total of 583 single-nucleotide polymorphisms (SNPs) based on a previous genome-wide association study (GWAS) of sleep behaviors were selected for the GRS of sleep index [21]. For the GRS of AAs, there is no relevant GWAS of KDM-biological age or PhenoAge for use. However, since the epigenetic clock based on DNA methylation age has been highly associated with BAs [19], we used 11 SNPs reported in a GWAS of epigenetic ages to create the GRS for both BAs/AAs [22]. Each SNP was recoded as 0, 1, or 2 according to the number of risk alleles and missing SNP values of individual were imputed by mean. The unweighted GRSs were directly summed up and the weighted GRSs were calculated by summing up after multiplying with its effect value, then divided half of the total of effect size. There is no overlapped SNPs for both GRSs and the GRS of BAs was highly correlated with both forms of BAs/AAs. The detailed SNPs were demonstrated in Table S1. The unweighted GRSs were employed in the primary analyses and the weighted ones were used in the sensitivity analyses.

### Statistical analysis

We first examined the associations of each sleep behavior and the sleep index with both forms of AAs using mixed-effect linear regression models in which the AAs were the outcomes. We adjusted for covariates described in the previous section and additionally controlled for the examination center in the model as a random effect to account for the potential residual bias from examinations. Then, we performed one-stage MR analyses to explore the causal relationships between sleep and BAs in two scenarios. The GRS of sleep index was used to predict sleep index genetically and to study the effect of sleep on BAs. The GRS of BA was used to predict AAs genetically and to study the effects of the two AAs on sleep quality. The statistically significant genetic predicted effect in either scenario that was in the same direction as the observed effect in the primary model would indicate the plausibility of a causal effect. Corresponding dose-response curves between sleep index and the two BAs were further assessed by restricted cubic spline regression [23]. Models were adjusted for the previously described covariates and the sleep index = 1, 3, and 5 were selected as knots.

Based on the results of MR analyses, the next part of our study would test whether (a) sleep quality could mediate or modify the association between elevated air pollution levels and accelerated AAs (i.e. if declined sleep index → the increased AAs), or (b) AAs could mediate or modify the association between elevated air pollution levels and declined sleep quality (i.e. if the increased AAs → declined sleep index). The possibility of the mediator role of the factors of interest would be examined by comparing the estimates of air pollutants in models with air pollutants only with the ones in models with the mutual adjustment of both air pollutants and the potential mediator. If a robust mediation effect was identified, the mediation proportion would be estimated by the mediation analysis using the SAS function ‘PROC CAUSALMED’. Meanwhile, the possible modifying effects of the factors of interest would be examined by an interaction term of air pollution and the factor. If the interaction terms were statistically significant and hence suggested a likely modifying effect, a subgroup analysis by sleep index category or binary AAs (by median) would be conducted to test the effects of air pollution on AAs or sleep quality. The effects of air pollutants in all models were demonstrated by per one interquartile range (IQR) increase in the concentrations. Corresponding dose-response curves and linearity of the relationships between air pollutants and BAs by sleep index category were subsequently assessed by restricted cubic spline regression if necessary. The 5^th^, 50^th^, and 95^th^ percentiles were selected as knots.

SAS version 9.4 TS1M5 (SAS Institute Inc., Cary, NC, USA) was used to conduct data cleaning and all analyses. A two-sided *p*-value of <0.05 was considered statistically significant.

## Results

### Participants’ characteristics and air pollution distributions

Table 1 presents the baseline characteristics of 363,886 study participants by sleep index category. Participants’ age (mean±standard deviation) was 56.5±8.1 years and most of them are white. About 35% and 55% participants were former and never smokers, respectively. Majority of them were with healthy physical activity and >10 years of education. Nearly half were with a healthy daily intake of alcohol. Only about 24% and 5% participants were with prevalent hypertension or diabetes diagnosed by doctors, respectively. Insomnia is the most frequent (∼75%) sleep disorder that the participants had and most participants (∼82%) could get up easily in the mornings. About 30% have a high sleep quality (sleep index = 5-6) and 15% have a low sleep quality (sleep index = 0-2). The high sleep quality group has lower BAs and higher AAs than the low sleep group, and the medium sleep quality group (sleep index = 3-4) has the BAs and AAs at the intermediate level in between. Both BAs were highly correlated with the chronological age and mutually correlated (Figure S1). The average concentrations of air pollutants were 9.96±1.05 μg/m^3^ (IQR=1.27) for PM_2.5_, 6.42±0.90 μg/m^3^ (IQR=0.79) for PM_coarse_, 19.23±2.01 μg/m^3^ (IQR=2.33) for PM_10_, 28.93±9.10 μg/m^3^ (IQR=10.80) for NO_2_, and 43.58±15.47 μg/m^3^ (IQR=16.44) for NOx. Levels of all air pollutants were highly correlated (Table S2, all *p*-values <0.001).

### Associations of sleep with biological ages

We first examined the associations of both forms of AAs with each of the six sleep behaviors (Table 2). After controlling for all potential covariates, we found that four and five sleep behaviors were negatively associated with KDM-biological age acceleration and PhenoAge acceleration, respectively. Early chronotype, normal sleep duration, and never or rarely have insomnia were significantly associated with both AAs. For instance, normal sleep duration was associated with 0.248 and 0.220-year decreases in KDM-biological age acceleration and PhenoAge acceleration, respectively. And never or rarely have insomnia was associated with 0.082 and 0.041-year decreases in KDM-biological age acceleration and PhenoAge acceleration, respectively. KDM-biological age acceleration was additionally associated with self-reported snoring, and PhenoAge acceleration was additionally related to frequent daytime sleepiness and difficulties in getting up in mornings. We further observed significant negative associations between the continuous sleep index and both AAs (Table 2 and Figure 1a-b). One unit increase in the sleep index was associated with 0.105 and 0.125-year decreases in KDM-biological age acceleration and PhenoAge acceleration, respectively. The AAs of high sleep quality group were 0.341 (KDM) and 0.467-year (PhenoAge) lower than the low sleep quality group, and the changes in both AAs of the medium sleep quality group were at the intermediate levels. As illustrated in Figure 1c and 1d, monotonic negative dose-response relationships between continuous sleep index and both AAs were observed.

**Figure 1.**
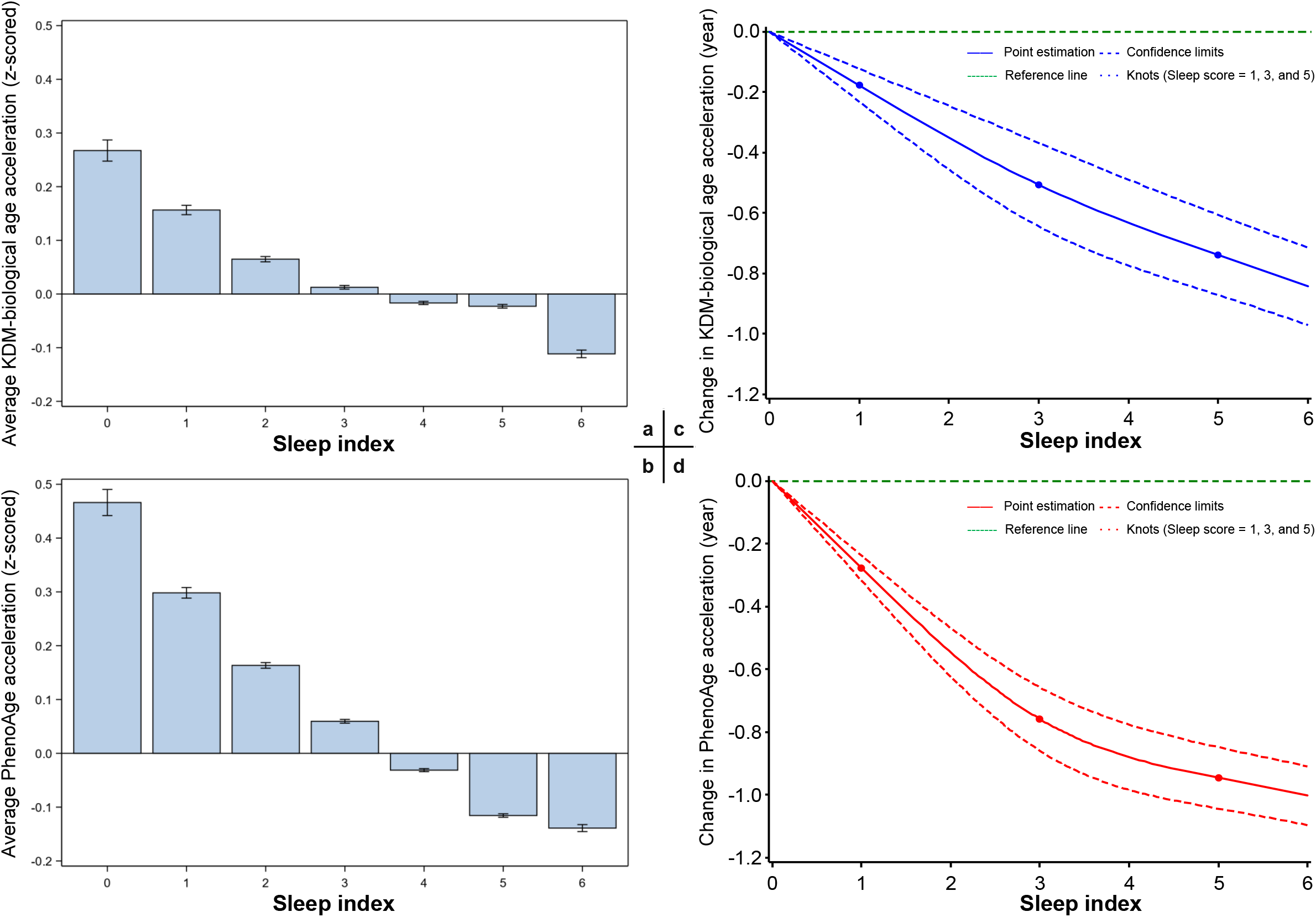
Distributions of age accelerations by sleep index and best-fitting dose-response curves

Furthermore, to investigate the causal associations between sleep and BAs, we conducted two MR analyses in two directions. As demonstrated in Table 3 with unweighted GRSs for sleep index and BAs, we observed significant negative associations between genetic-predicted sleep index and both AAs. However, the genetic-predicted KDM-biological age acceleration was positively associated with sleep index, and the negative association between genetic-predicted PhenoAge acceleration and sleep index was not statistically significant. Sensitivity analyses with weighted GRSs demonstrated similar trends for each scenario (Table S3). These results indicated that sleep quality was more likely to be the determinant of the change in AAs rather than a consequence.

### Joint effects of air pollution and sleep on biological ages

Given it was more plausible that accelerated AAs were the consequences of impaired sleep quality, along with the known association between air pollution and biological aging, we validated the associations of the five air pollutants with each BA and explored the hypothesis that whether sleep index could mediate the effects of air pollutants on BAs. As shown in Table 4, PM_2.5_ and NO_2_ were the two air pollutants that were significantly associated with both AAs in models without the adjustment of sleep index. An IQR increase in PM_2.5_ level was associated with 0.061 and 0.045-year higher KDM-biological age acceleration and PhenoAge acceleration (*p*-values <0.0001), respectively. The same change in NO_2_ level was associated with 0.098 and 0.096-year higher KDM-biological age acceleration and PhenoAge acceleration (*p*-values <0.0001), respectively. Their effects and *p*-values were essentially unchanged in models which additionally controlled for sleep index. Meanwhile, the effects of sleep index on both AAs were also very slightly altered in this mutual adjustment model. Therefore, we suggest that air pollution and sleep quality may be independently associated with BAs. To understand their joint effects explicitly, we classified the participants based on binary air pollution levels (by median) and sleep index category, and then observed clear and stepwise increasing trends of the joint effects of both factors on the two AAs (Figure 2). Particularly, compared to the group with high sleep quality and lower exposure to PM_2.5_, people with low sleep quality and higher exposure had a 0.405- and 0.543-year higher KDM-biological age acceleration and PhenoAge acceleration (*p*-values <0.0001), respectively (Table S4).

**Figure 2.**
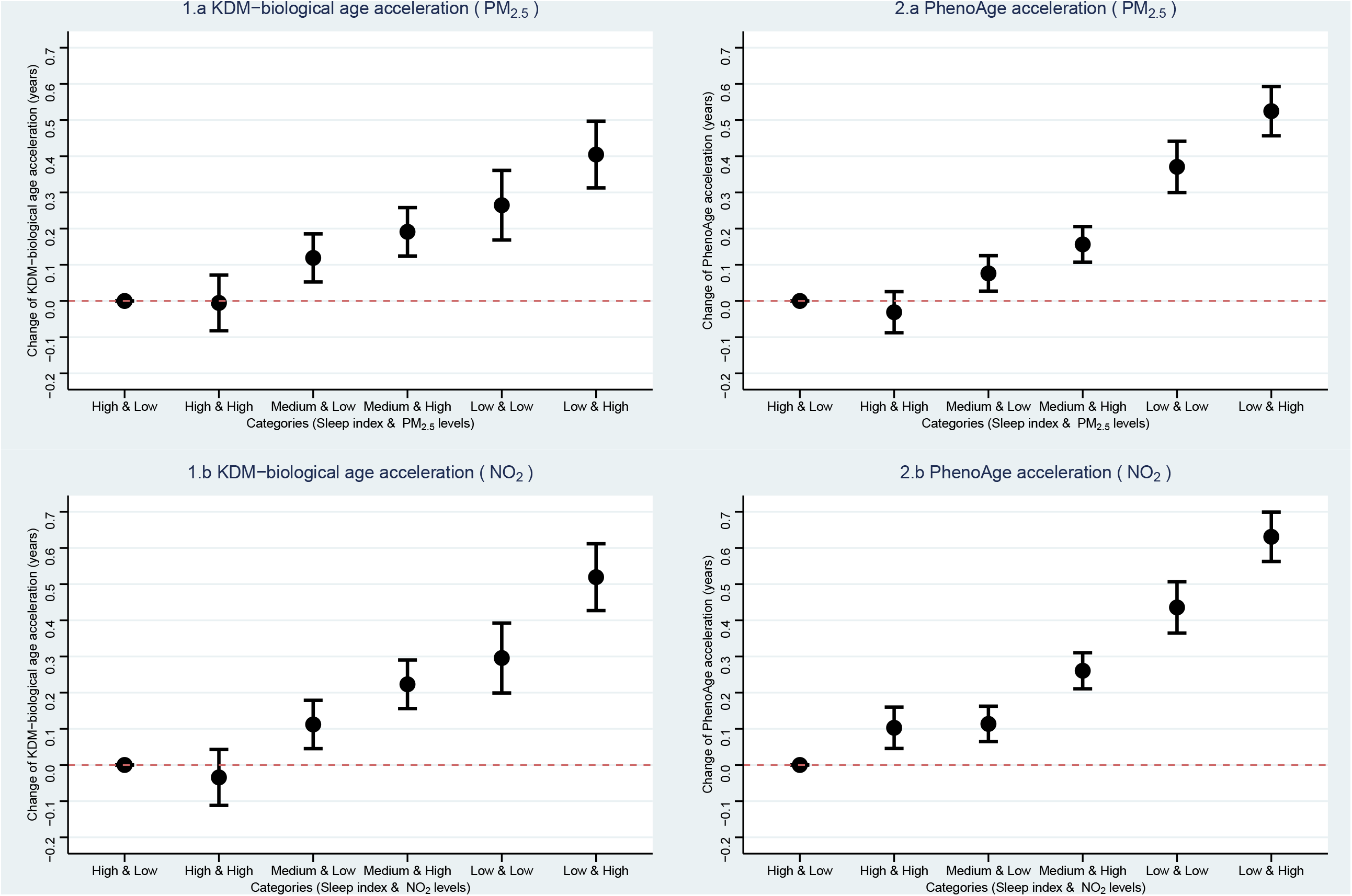
Joint associations of sleep index and air pollution levels with biological age accelerations

### Modifying effects of sleep quality

According to the robust joint effects of air pollution and sleep on BAs, we additionally explored the potential modifying effect of sleep quality on the associations between air pollution and BAs since sleep is a modifiable factor that could be easily intervened by human behaviors. As shown in Figure 3 and Table S5, we observed a robust modifying effect of sleep quality on the air pollution-AA associations (*p*-values of interaction terms <0.01). Particularly, an IQR increase in PM_2.5_ concentration was associated with 0.011-, 0.047-, and 0.078-year increase in PhenoAge acceleration among people with high, medium, and low sleep quality, respectively. The same increase in NO_2_ concentration was associated with 0.049-, 0.100-, and 0.138-year increase in PhenoAge acceleration among people with different sleep qualities. We further evaluated the linearities and dose-response relationships of the PM_2.5_ and NO_2_ with both AAs. Non-linear relationships with respect to each air pollutant were observed (Figure 4 and Figure 5). Modifying effects of sleep quality on the changes in the KDM-biological age acceleration could be distinguish under relatively lower PM_2.5_ (∼5 μg/m^3^, Figure 4a) and NO_2_ levels (∼20 μg/m^3^, Figure 5a) than corresponding modifying effects of sleep quality on the changes of PhenoAge acceleration (PM_2.5_: ∼10 μg/m^3^, Figure 4b; NO_2_: ∼40 μg/m^3^, Figure 5b).

**Figure 3.**
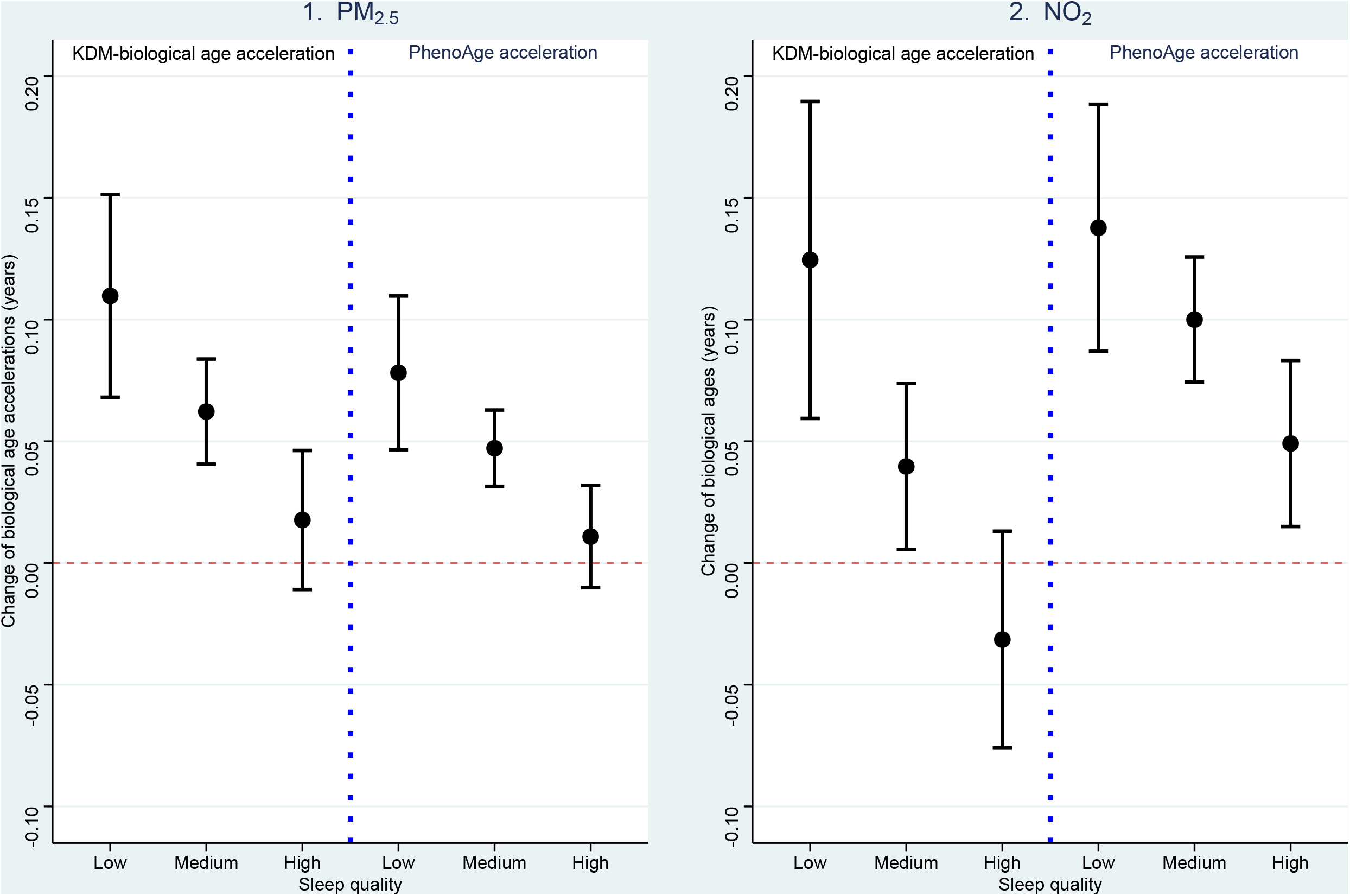
Associations of air pollution levels with biological age accelerations by sleep quality

**Figure 4.**
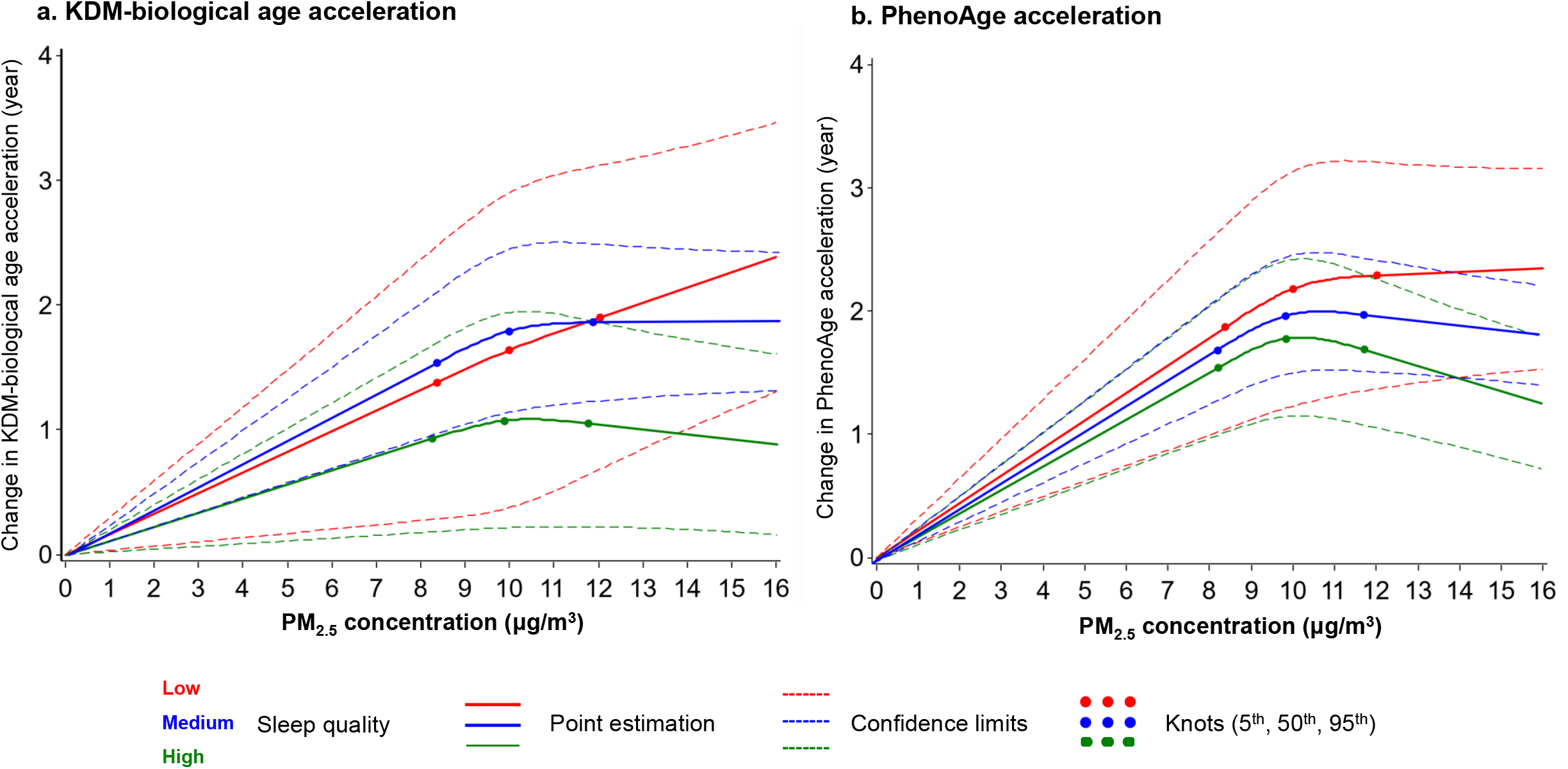
Best-fitting models for the relationships of PM_2.5_ exposure with accelerations of two biological ages, by sleep quality

**Figure 5.**
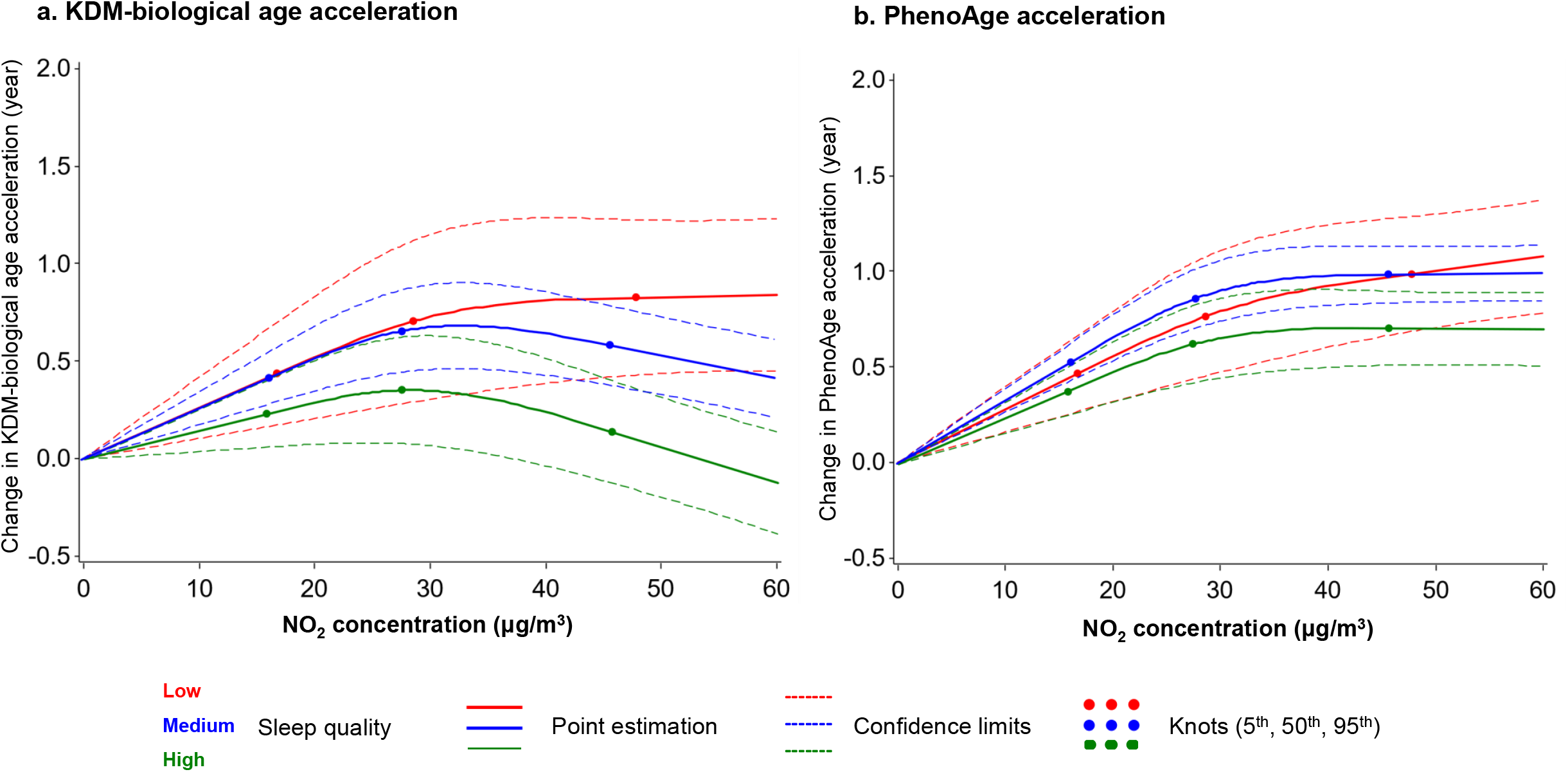
Best-fitting models for the relationships of NO_2_ exposure with the accelerations of biological ages, by sleep quality

## Discussion

In this large cohort of middle and elderly-aged adults, we demonstrated that worsening sleep quality could accelerated biological aging with BAs estimated by two well-established algorithms and a comprehensive sleep index combining the impacts of six major sleep disorders. We further observed that people with low sleep quality and higher exposure to PM_2.5_ or NO_2_ had the highest AAs. Sleep quality could also modify the associations of elevated PM_2.5_ and NO_2_ levels with accelerated aging. For instance, an IQR increase in PM_2.5_ concentration was associated with 0.011-, 0.047-, and 0.078-year increase in PhenoAge acceleration among people with high, medium, and low sleep index, respectively.

To date, our study is the first investigation demonstrating that impaired sleep quality could accelerate human aging in a large population with the state-of-art of causal interpretation. Our observed negative sleep-BA association has been partially suggested by previous studies. In Han et al.’s study, they reported a positive association of sleep duration of >8h per day with an increased BA estimated based on five phenotypes [11], which was also found in our study that abnormal sleep duration was associated with increased AAs. Additionally, Carroll et al. and Carskadon et al. separately linked insomnia and insufficient sleep duration with accelerated epigenetic clocks among older females [9], pregnant women [10], and freshmen [12], respectively. In line with these studies, we observed increasing patterns of AAs under worsening general sleep quality. Previous studies also provided much more marginal evidence that is consistent with our findings with respect to other aging biomarkers. For example, in a study of physicians, overnight on-call participants had lower baseline DNA repair gene expression and more DNA breaks than participants who did not work overnight [24]. And shortened telomere length, a well-known biomarker of cellular aging, was also found in relation to <6 h of sleep [25] and chronic poor sleep quality [26], respectively. Sun et al. also preliminarily found that poor sleep was associated with being frailty, another aging-related syndrome [13]. Taken together, these complex aging biomarkers including BAs provided an exciting new avenue for investigating the underlying biological mechanisms of the interplay between sleep and aging-related health outcomes.

More intriguingly, although aging is tied to sleep difficulties firmly, due to the relatively limited sample size and cross-sectional nature, previous studies were not able to address the causal associations. Our study answered this scientific question of interest by a causal inference test using the MR scheme. We found that it was more likely that sleep behaviors lead to the changes in BAs with statistically significant associations between genetic-predicted sleep index and both AAs. This suggests that sleep quality may affect the aging of organs and systems reflected by two BAs based on mainly blood biochemistry biomarkers, and indicates that improving modifiable aspects of sleep may help to lessen the adverse effects associated with biological aging and to attenuate the risks of aging-related diseases. Since BAs have been associated with cardiovascular health [27], our finding is in agreement with that sleep quality is a preventable risk factor of cardiovascular events, which have been well-established and recognized in large population-based studies and clinical trials [28]. However, our study did not fully exclude the possibility that aging could be one of the major determinants of sleep impairment as we also found a negative correlation between genetic-predicted PhenoAge acceleration and sleep in MR analyses, albeit not significant. Given multiple factors including environmental, social, lifestyle, medical, and psychiatric conditions may contribute to aging acceleration [4], the genetic-predicted PhenoAge acceleration may not be able to capture the contributions from non-genetic backgrounds to aging. The GRS of BAs/AAs may be another cause, because we did not find the SNPs that are directly associated with both BAs but with epigenetic clocks, this GRS may not an ideal instrumental variable for both BAs though shared high correlations with both. Both reasons may contribute to the null causal effect of aging on sleep impairment in our study.

Furthermore, literatures have established the adverse impact of air pollution on sleep quality in different perspectives [15], such as the risks of breathing problems, insomnia, sleep efficiency, and overall sleep quality, as well as on aging in different populations [14]. With the previously determined causal association between sleep and BAs in the first step, we expected to find a robust mediation effect of sleep on the air pollution-BA relationship. However, we observed very limited changes in the effects of different air pollutants on BAs in models mutually adjusting for sleep index, which suggests that sleep does not play a major role in mediating the effects of air pollution on the two BAs. Since sleep is predominantly related to cognitive health and brain aging [3], especially the structural and physiological changes that occur in the brain, such limited mediation effects could be explained by the minimal capacity of the two BAs in measuring aging-related neurological changes. The clinical traits implemented in the constructions of the two BAs were not specifically associated with such abnormal alterations related to the central neuron system. Instead, we observed prominent modifying effects of sleep on the air pollution-BA relationship, which suggests that a healthy and adequate sleep may help attenuate the adverse aging effects of air pollution on non-neurological systems. Also, we noted response curves with different features of the PM_2.5_ and NO_2_ with both AAs. Because KDM-biological age is more related to the capacity and function of systems and organs [29], and PhenoAge is skewed to predict the mortality risk of humans [7]. Such various features of the modifying effects of sleep on the AAs suggest that sleep may help lessen the detrimental impacts of PM_2.5_ and NO_2_ on body functions at a lower exposure level and could also attenuate the lethal effects of the two pollutants on mortality when they reached a higher level.

Regarding the underlying biological mechanisms under the interesting modifying effects of sleep, one of the plausible explanations is the dampen of oxidative stress and inflammation during healthy and adequate sleep. First, air pollutants can induce oxidative stress, the ability to respond to which has been identified as a key determinant of biological aging [14]. Sleep may help with the anti-oxidative mechanism by removing reactive oxygen species resulting from air pollution insults [30]. For instance, it has been suggested that sleep reduces the accumulation of free radicals accumulated during wakefulness [31]. Beyond this, sleep may also help enhance immune defenses and stabilize the dysregulation of inflammatory responses under air pollution exposure. Previous studies have suggested that sleep impairment is associated with increased serum levels of C-reactive protein, interleukin (IL)-1, IL-6, IL-17, tumor necrosis factor α (TNF-α), and nuclear factor-kappa B (NF-κB) in addition to alteration of numbers and activity of macrophages and natural killer cells [30, 32]. The aberrant changes in such systematic inflammation biomarkers have been found to be involved in the impact of air pollution on the integrity of organs and the development of aging-related diseases [33].

The major strengths of this study include the large sample size, the detailed records of sleep behaviors, and rich phenotype and biochemistry data for the biological age estimation. Several limitations are notable when interpreting the results. First, UK Biobank is a volunteer cohort, and participants are likely healthier than the general population, which may limit the effect of sleep on BAs in our analysis as their AAs are expected to be lower than the general population theoretically. Furthermore, the measurement bias of air pollution must be noted. The air pollution data we used were mostly only a single measurement of the annual average outdoor air pollution level in 2010 since home addresses of the participants are unavailable during follow-up. Because the initial assessment visit of UK Biobank was from 2006 to 2010, we were unable to determine the lag or short-term (<1 month) effects of air pollution on both sleep and BAs. Also, as most individuals spend a large amount of time indoors, individual exposure to all forms of air pollution may differ from that indicated by the ambient outdoor levels we used. Additionally, self-reported sleep data was used in our analyses, misclassification of sleep behaviors was therefore inevitable. However, such bias may attenuate our findings toward the null and underestimate the effects we observed.

Meanwhile, our sleep index dichotomized six sleep behaviors for simplicity but did not include all sleep behaviors or take the changes of sleep behaviors before and after the survey into consideration, which may cause residual bias to some extents. Last, participants in this study were mostly of European descent, which limits the generalization of the results to other ethnic groups.

In conclusion, our study is the first identifying the accelerating effect of poor sleep quality on biological aging. With this premise, we further found that sleep and air pollution were independently associated with biological aging, and sleep quality may modify the aging effects of air pollution. These findings not only provide solid evidence supporting sleep as an aging contributor but also underscore the importance of high-quality sleep as an intervention approach to mitigate the negative impact of air pollution on human aging. Nevertheless, aging is associated with a myriad of changes in psychological, social, spiritual, financial, and lifestyle over the lifetime [4]. Further longitudinal studies with a more detailed landscape of aging are highly warranted to validate our findings and further determine the underlying biological mechanisms.

## Acknowledgments

Data are available in a public, open access repository. This research has been conducted using the UK Biobank Resource under Application Number 44430. The UK Biobank data are available on application to the UK Biobank (www.ukbiobank.ac.uk/). Dr. Xu Gao was supported by grants from the Peking University Start-up Grant (BMU2021YJ044). We thank Dr. Chen Chen for the language assistance.

## Author contributions

Xu Gao conceptualized the study, conducted data clean, estimated biological ages, and draft and reviewed the manuscript; Ninghao Huang coordinated the data collection, reviewed and revised the manuscript; Tao Huang coordinated and supervised the data collection, and critically reviewed and revised the manuscript;

## Supplementary Materials

**Table S1** Characteristics of genetic variants associated with sleep behaviors and biological ages in UK Biobank

**Table S2** Distributions and correlation matrix of the five air pollutants in UK Biobank (Pearson correlation)

**Table S3** Causal associations between sleep index and biological age accelerations with weighted genetic risk scores

**Table S4** Joint associations of sleep index and air pollution levels with biological age accelerations

**Table S5** Associations of air pollutants with biological age accelerations by sleep quality

**Figure S1.**
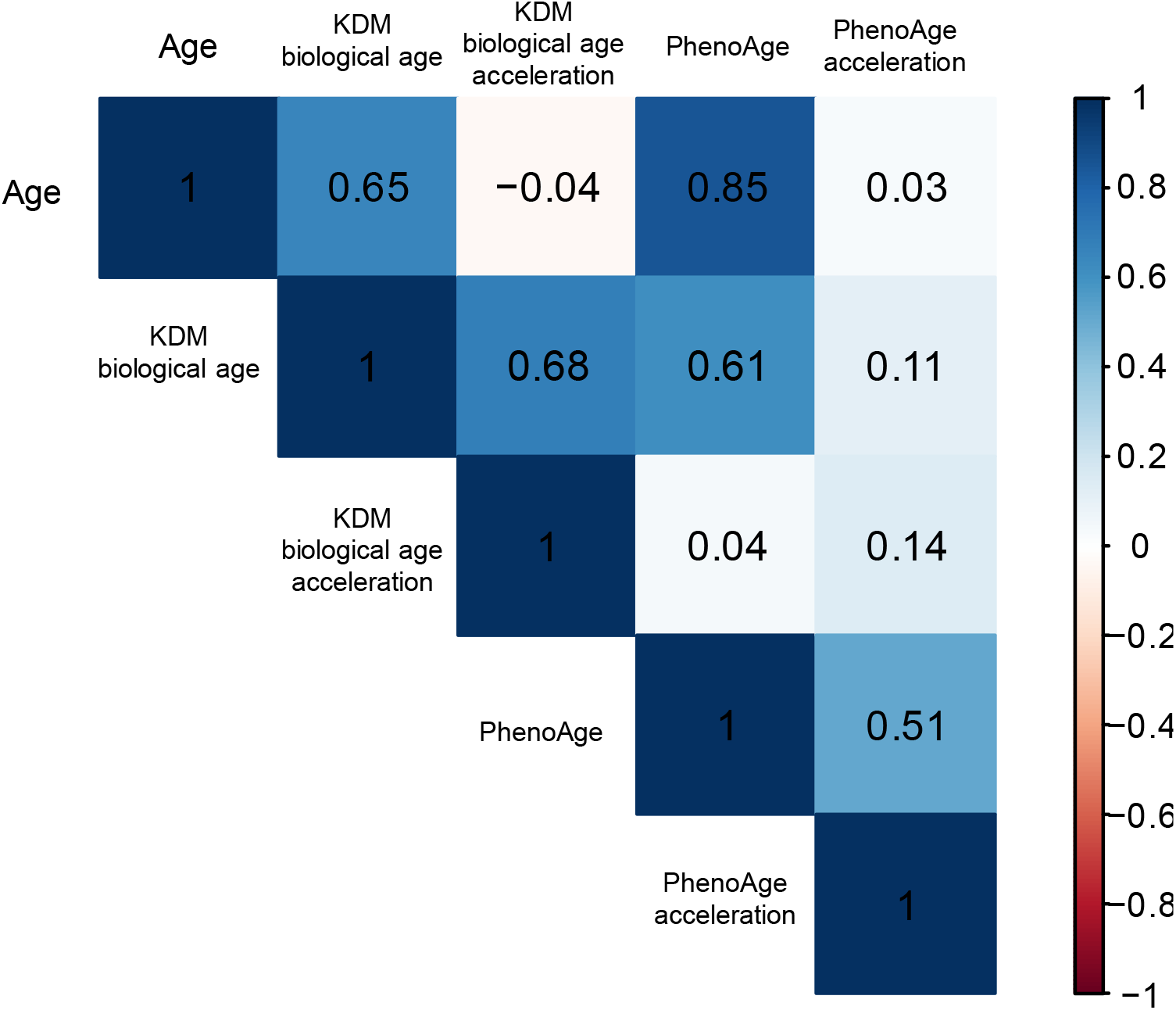
Correlation matrix of age and biological ages in UK biobank

